# *Bacillus safensis* FO-36b and *Bacillus pumilus* SAFR-032: A Whole Genome Comparison of Two Spacecraft Assembly Facility Isolates

**DOI:** 10.1101/283937

**Authors:** Madhan R Tirumalai, Victor G. Stepanov, Andrea Wünsche, Saied Montazari, Racquel O. Gonzalez, Kasturi Venkateswaran, George E. Fox

## Background

Microbial persistence in built environments such as spacecraft cleanroom facilities [1–3] is often characterized by their unusual resistances to different physical and chemical factors [1, 4–7]. Consistently stringent cleanroom protocols under planetary protection guidelines over several decades [1, 8–12], have created a special habitat for multi-resistant bacteria, many of which have been isolated and identified [13–19]. The potential of many of these isolates to possibly survive interplanetary transfer [2, 20–24] raises concern of potential forward and backward bacterial contamination. Understanding the survival mechanisms employed by these organisms is the key to controlling their impact on exobiology missions. In addition, their occurrence in the closed environments of the International Space Station, (ISS), could possibly impact the living conditions there as well [1–3, 25–27].

Two of the most studied organisms in the specialized econiches of spacecraft assembly facilities and the ISS are *B. safensis* FO-36b^T^ [28] (referred to as FO-36b henceforth) and *B. pumilus* SAFR-032 [16] (referred to as SAFR-032). These organisms are representative strains of the endospore producing *Bacillus* sp.[13, 16, 29–33]. Both strains produce spores that exhibit unusual levels of resistance to peroxide and UV radiation [24, 29, 34] that far exceed that of the dosimetric *B. subtilis* type strain (*B. subtilis subsp. subtilis str.* 168, referred to as BSU) [35]. A third strain, *B. safensis* MERTA-8-2 (referred to as MERTA), was initially isolated from the Mars Odyssey Spacecraft and associated facilities at the Jet Propulsion Laboratory and later also found on the Mars Explorer Rover (MER) before its launch in 2004. It has been reported that this strain actually grows better on the ISS than on Earth [36]. However, the resistance properties of its spores have not been directly tested. A recent phylogenetic study of 24 *B. pumilus* and *B. safensis* strains, found FO-36b, and MERTA clustered together in a distinct group of *B. safensis* strains [37].

Previously a draft genome of FO-36b with as many as 408 contigs (https://www.hgsc.bcm.edu/microbiome/bacillus-pumilus-f036b) was compared to SAFR-032 and the type strain *B. pumilus* ATCC7061^T^ [38, 39] (referred to as ATCC7061). This comparison identified several genes and a mobile genetic element in SAFR-032 that may be associated with the elevated resistance [39]. Since this previous study was completed, minor corrections to the SAFR-032 gene order were made and the annotation was updated [40]. In addition, a draft genome of MERTA was reported [41]. Herein, we now report a complete genomic sequence for FO-36b and the results of a detailed comparison of these four genomes.

## Methods

### Sequencing of the *Bacillus safensis* FO-36b genome

5μg of purified genomic DNA of FO-36b was digested with NEBNext dsDNA Fragmentase (New England Biolabs, Ipswich, MA) yielding dsDNA fragments in a size range of 50 bp up to 1000 bp. The fragments were fractionated on a 2% agarose gel, and those with the length from 300 bp to 350 bp were isolated as described [42]. The dsDNA fragments were converted to a shotgun DNA library using the TruSeq PCR-Free DNA Sample Preparation Kit LT (Illumina, San Diego, CA) according to the manufacturer’s instructions. Sequencing was performed on the Illumina HiSeq 2500 sequencer at the University of Arizona Genetic Core Facility (Tucson, AZ). A total of 10,812,117 pairs of 100 base-long reads with average Phred quality of 34.92/base were collected. The reads were processed with Sickle 1.33 [43] and Trimmomatic 0.32 [44] was used to remove seven 3′-terminal low-quality bases, and to filter out the reads with average Phred quality below 16/base as well as reads containing unidentified nucleotides. Overall, 9,047,105 read pairs and 1,435,623 orphaned single reads with a total of 1,816,274,469 nucleotides were retained after the filtration step. The reads were assembled using the Abyss 1.5.2 *de novo* assembler [45] with the *kmer* parameter set at 64. The assembly consisted of 22 contigs with a total length of 3,753,329 bp. The average contig length was 170,605 bp (ranging from 352 to 991,464 bp), with an N50 contig length equal to 901,865 bp. Data from two previous FO-36b draft genomes (https://www.hgsc.bcm.edu/microbiome/bacillus-pumilus-f036b; and https://www.ncbi.nlm.nih.gov/biosample/SAMN02746691) did not provide the additional information needed to order the 22 remaining contigs.

Instead, connections between the contigs were obtained by systematic PCR screening using LongAmp *Taq* DNA polymerase (New England Biolabs, Ipswich, MA) and near-terminal outward-facing primers. The amplicons were gel purified and sequenced by the Sanger method at SeqWright, Inc (Houston, TX). This allowed closure of all the gaps between the contigs. The complete FO-36b genome sequence comprises 3.77 Mb and has G+C content of 41.74%.

### *B. safensis* FO-36b genome annotation

The FO-36b genome was annotated using the NCBI’s Prokaryotic Genome Annotation Pipeline [46]. 3850 ORFs and 40 non-coding RNAs and riboswitches were predicted and the results were deposited in Genbank under accession number CP010405.

### Genomes used in comparisons

The recently updated complete sequence of the SAFR-032 genome was obtained from NCBI (CP000813.4). The draft genomes of ATCC7061^T^ (Refseq accession no: NZ_ABRX00000000.1), consisting of 16 contigs and MERTA consisting of 14 contigs (Refseq accession no: GCF_000972825.1) were obtained from the public databases of the National Center for Biotechnology Information (NCBI). Several additional *B. safensis* and *B. pumilus* draft genomes from various sources have also been deposited in the NCBI database in recent years. However, these genomes get excluded when performing a global Genbank Blast (NT) analysis. To avoid this potential problem, these additional draft genomes were separately retrieved from the Genbank repository (*B. pumilus* genomes, https://www.ncbi.nlm.nih.gov/genome/genomes/440; *B. safensis* genomes, https://www.ncbi.nlm.nih.gov/genome/genomes/13476) and locally integrated into the Genbank NT database. The resulting local database allowed inclusion of these genomes in subsequent Blast (NT) studies. Overall, the analysis involved 65 *B. pumilus* and *B. safensis* genomes (including the FO-36b, MERTA, SAFR-032 and ATCC7061 genomes). The names of the genomes used are given in Additional file 1: Table S1

### BLAST studies

Individual gene and protein sequences from the FO-36b genome, were blasted against each other as well as against the genomes of SAFR-032, MERTA and ATCC7061 using the standalone version of NCBI’s BLAST program [47]. The comprehensive search included blastN and blastX for the nucleotide sequences and blastP for the protein sequences. Additionally, global blast was performed on the sequences against the updated NR/NT databases downloaded from the NCBI on the Opuntia Cluster at the Center of Advanced Computing and Data Systems at the University of Houston.

Genes with BLAST results in which the best hit had an e-value greater than (an arbitrary) 0.0001 were considered absent from the target genome, while those with BLAST e-values below e-10 were considered to be matches. Genes with e-values between e-20 and 0.0001 were further analyzed by aligning the sequence of the entire gene neighborhood with the corresponding region in the other genomes to ascertain/verify the BLAST results as well as to look for unusual features in the sequence. Gene/protein sequence alignments were performed using Bioedit (http://www.mbio.ncsu.edu/BioEdit/bioedit.html).

### Phage analysis

The online tool PHAST [48, 49] was used to predict and annotate potential phage elements in the genomes. Comparative analysis of the respective homologs on the other genomes, were performed to map the respective corresponding phage regions on the other genomes.

### Whole Genome Phylogenetic Analysis (WGPA) and Genome-Genome Distance Studies (GGDC)

In order to obtain an overall view of relationships among the various genomes, we used seven additional genomes thereby forming a complete set of 72 strains. Overall, the genomes included 65 *B. pumilus* and *B. safensis* genomes (including those of FO-36b, MERTA, SAFR-032 and ATCC7061), four representative strains from the *B. altitudinis* complex, *viz*., *B. aerophilus* C772, *B. altitudinis* 41KF2b, *B. cellulasensis* NIO-1130(T), and, *B. stratosphericus* LAMA 585.

The genomes of *Geobacillus kaustophilus*, and *B. subtilis* served as outliers in the *Firmicutes* group, while the genome of Gram-negative *E. coli* MG1655, served as a non-*Firmicutes* outlier.

A whole-genome-based phylogenetic analysis was conducted using the latest version of the Genome-BLAST Distance Phylogeny (GBDP) method [50] as previously described [51]. Briefly, BLAST+ [52] was used as a local alignment tool and distance calculations were done under recommended settings (greedy-with-trimming algorithm, formula D5, e-value filter 10e-8). 100 pseudo-bootstrap replicates were assessed under the same settings each. Finally, a balanced minimum evolution tree was inferred using FastME v2.1.4 with SPR post processing [53]. Replicate trees were reconstructed in the same way and branch support was subsequently mapped onto the tree. The final tree was rooted at the midpoint [54]. The genomes were also compared using the in-silico genome-to-genome comparison method, for genome-based species delineation and genome-based subspecies delineation based on intergenomic distance calculation [50, 55].

In order to confirm the reasonableness of these results, a separate analysis was conducted using DNA gyrase A (*gyrA*), which has often been used for single gene phylogenetic studies [28, 56–60]. *gyrA* is preferable to 16S rRNA in this case, because many of the 16S rRNAs are too similar [61]

The *gyrA* sequences were bioinformatically isolated from all 72 genomes and aligned using Bioedit ((http://www.mbio.ncsu.edu/BioEdit/bioedit.html), ClustalW, and MEGA [62, 63] with MUSCLE. Maximum Likelihood, Neighbor-Joining and Minimum Evolution trees were built using MEGA. The Maximum Likelihood tree was built using the Tamura-Nei model [64]. The tree with the highest log likelihood (-18473.7156) was used. Initial tree(s) for the heuristic search were obtained automatically by applying the Neighbor-Join and BioNJ algorithms to a matrix of pairwise distances estimated using the Maximum Composite Likelihood (MCL) approach. The topology with superior log likelihood value was selected.

A Minimum Evolution (ME) Tree was built using the method described by Rzhetsky and Nei (1992) [65]. The ME tree was searched using the Close-Neighbor-Interchange (CNI) algorithm [66] at a search level of 1. The Neighbor-Joining (NJ) Tree was built using the method described by Saitou and Nei (1987) [67].

For both the ME and NJ trees, the optimal tree(s) with the sum of branch length = 1.62873358 was derived. The evolutionary distances were computed using the Maximum Composite Likelihood method [68] and are in the units of the number of base substitutions per site.

The analysis involved 72 nucleotide sequences. Codon positions included were 1st+2nd+3rd+Noncoding. All positions containing gaps and missing data were eliminated. There were a total of 2424 positions in the final dataset. Evolutionary analyses were conducted in MEGA6 [69].

The Mauve alignment [70] program was used to align the previous draft FO-36b sequence (GCA_000691165.1 / ASJD00000000) with the current updated sequence (CP010405).

### Screening Genomes for Antibiotic resistance genes

A global analysis of each of the four genomes was performed to identify possible antibiotic resistance loci. This was done using the reference sequences of the Comprehensive Antibiotic Resistance Database (“CARD”) [71], In addition a search for potential ‘resistome(s)’ was undertaken using the Resistance Gene Identifier feature of the CARD database for the four genomes.

## Results

### Unique and characteristic genes

Genes are considered to be characteristic if they are present in FO-36b, but absent in the other three organisms examined here. Unique genes are those that are not only absent in the other three genomes, but have not yet been found in any other genome. 307 ORFs found in FO-36b are not shared by SAFR-032. Sixty five of these ORFs did not have homologs in the genomes of ATCC7061 or MERTA and are therefore considered characteristic (Table 1). Although most are open reading frames that code for hypothetical proteins, six genes suggest that FO-36b has a CRISPR system. The likely presence of a CRISPR system is shared by 5 other *B. safensis* genomes and 8 other *B. pumilus* genomes (Additional file 2: Table S2). Among the 49 hypothetical protein coding ORFs, 26 are predicted to be part of phage element(s).

**Table 1:**
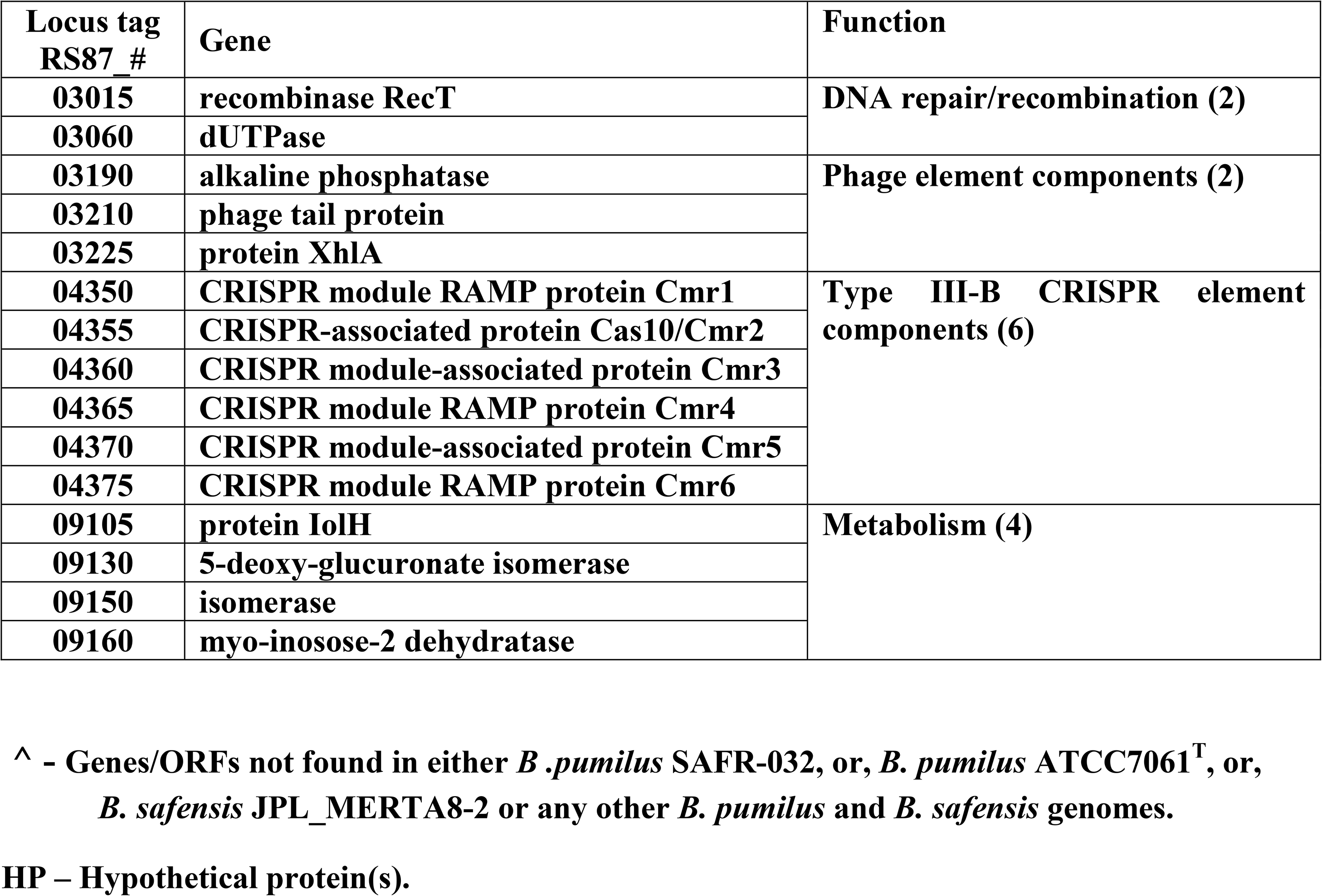
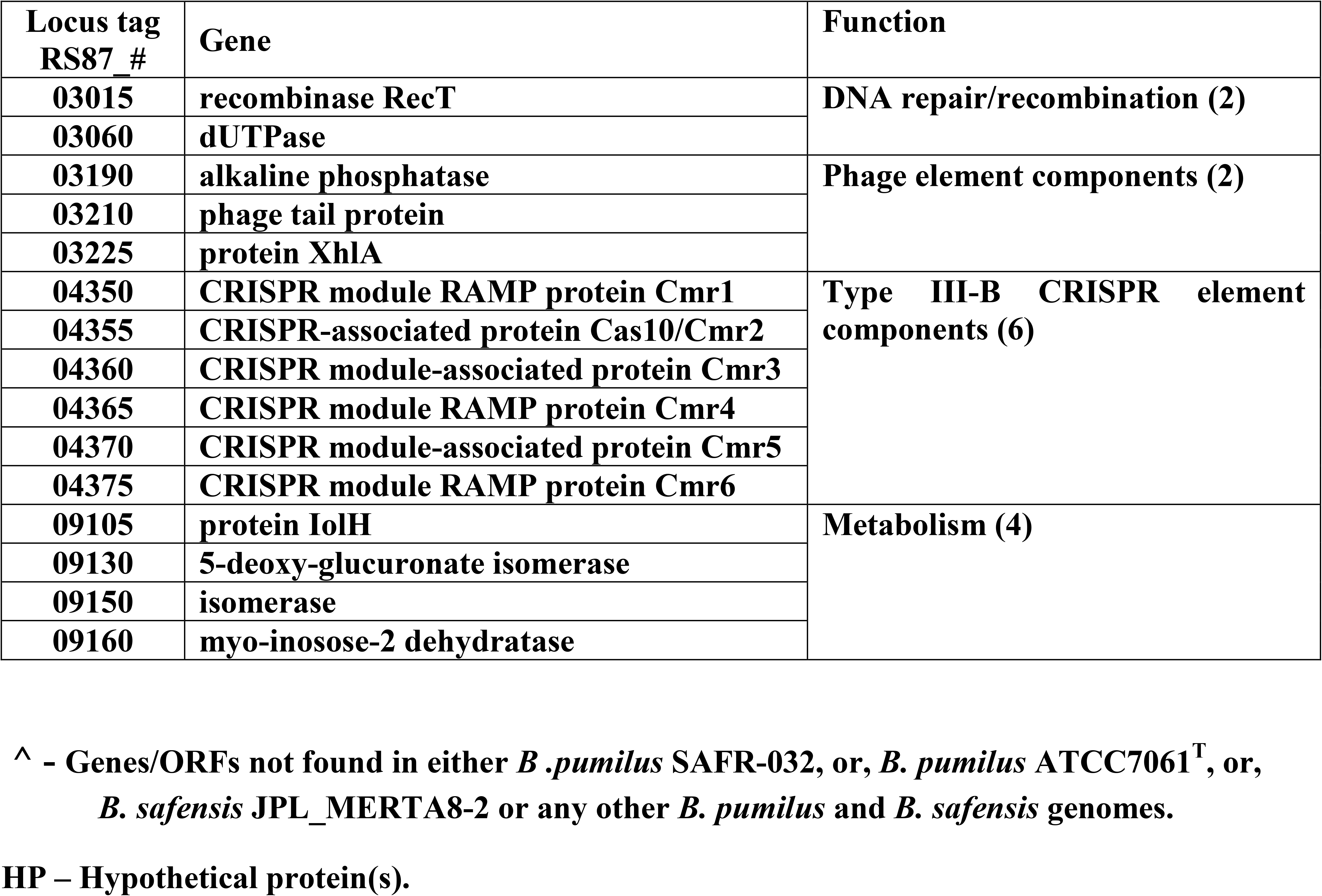
List of *B. safensis* FO-36b (characteristic) genes not shared by *B. pumilus* SAFR-032, *B. pumilus* ATCC7061^T^ and *B. safensis* JPL_MERTA8-2.

The analysis was extended to all available genomes of *B. safensis* (https://www.ncbi.nlm.nih.gov/genome/genomes/13476) and *B. pumilus* (https://www.ncbi.nlm.nih.gov/genome/genomes/440). Nine ORFs/genes classified as FO-36b characteristic are absent from all the *B. safensis* and *B. pumilus* genomes available in the NCBI database. These nine genes are totally unique to FO-36b with no homologs in the entire NR/NT databases (Table 2). Four of these are part of predicted phage elements. In addition, there are four genes with fewer than five homologs found in other *B. pumilus*/ *B. safensis* genomes (Table 3). Overall 217 SAFR-032 ORFs are not shared by *B. safensis* FO-36b. Sixty three of the 65 FO-36b characteristic ORFs are absent in 28 of the 61 total *B. safensis, B. pumilus*, and *Bacillus sp*. WP8 genomes. 18 are absent in all the *B. safensis* genomes, while 15 are not found in any of the *B. pumilus* genomes (Additional file 3: Table S3).

**Table 2:** FO-36b unique genes (hypothetical proteins)

| **Locus tag RS87_#** |
| --- |
| **03140\*** |
| **09820** |
| **12770** |
| **14110** |
| **14145** |
| **14150** |
| **14155\*** |
| **14285\*** |
| **14310\*** |

**\* Genes that are part of phage elements**

**Table 3:** FO-36b genes (hypothetical proteins) with fewer than 5 homologs.

| **Locus tag RS87_#** |  |
| --- | --- |
| **03030\*** | **only four homologs in**<br>***B.safensis* U41 (GCA_001938685.1),**<br>***B.safensis* U17-1 (GCA_001938705.1),**<br>***B.pumilus* CCMA-560 (GCA_000444805.1), and,**<br>***B.pumilus* strain 36R_ATNSAL (GCA_002744245.1).** |
| **03050\*** | **only one homolog in *B.pumilus* strain 36R_ATNSAL (GCA_002744245.1).** |
| **03110** | **only two homologs in**<br>***B.safensis* 7783 (GCA_002276315.1), and,**<br>***B.safensis* Bcs96 (GCA_002155005.1).** |
| **04125** | **only one homolog in *B.pumilus* PE09-72 (GCA_002174275.1).** |

**\* Genes that are part of phage elements**

### Phage insertions

The genome of FO-36b contains two phage insertions, namely the *Bacillus* bacteriophage SPP1 (NC_004166.2) insertion and the *Brevibacillus* phage Jimmer 1 (NC_029104.1) insertion. The SPP1 insertion, (Figure 1), consists of 62 genes (RS87_02955 to RS87_03255). Abbreviated versions are found in the MERTA strain (4 genes) and the ATCC7061 strain (3 genes), (Figure 1 and Figure 2). Portions of this element can also be detected in other *B. safensis/ B. pumilus* strains by sequence comparison.

**Figure 1.**
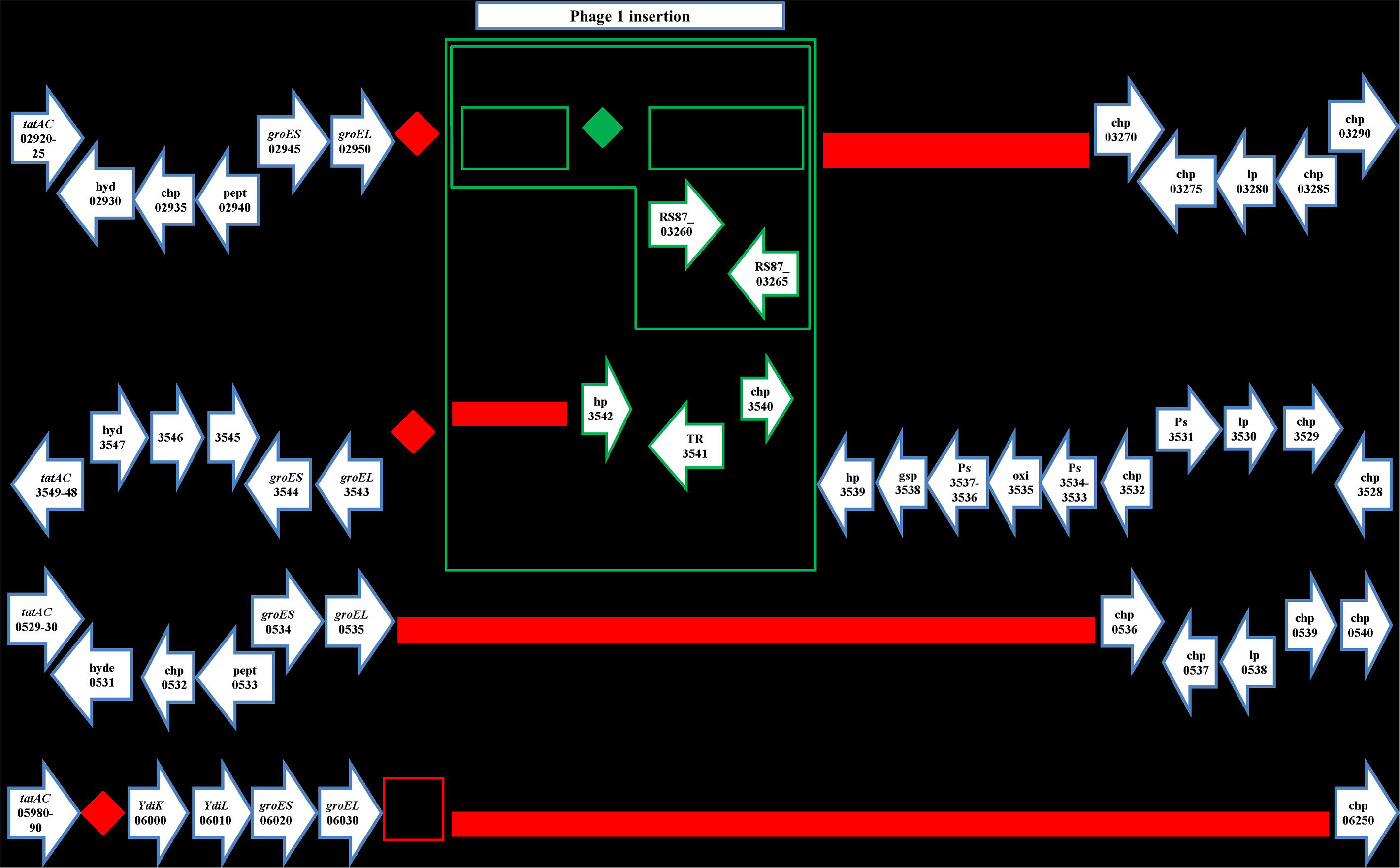
The *Bacillus* bacteriophage SPP1 (NC_004166) homologous region in the *B. safensis* FO-36b genome, as compared with the equivalent genomic regions of *B. pumilus* ATCC7061^T^, *B. pumilus* SAFR-032 and *B. subtilis subsp. subtilis str.* 168. The locus tag numbers are given inside the boxes/rectangles. Red diamonds denote absence of a single gene/homolog. Red rectangle denotes absence of a series/cluster of ORFs/genes. Green box encloses the phage insertion region. Green diamond denotes absence of a single gene/homolog within the phage. “hyd” = hydrolase, “chp” = conserved hypothetical protein, “pept” = peptidase, “hp” = hypothetical protein, “TR” = transcriptional regulator, “Ps”= pseudogene, “lp” = lipoprotein, “gsp” = group specific protein, “oxi” = oxidase.

**Figure 2.**
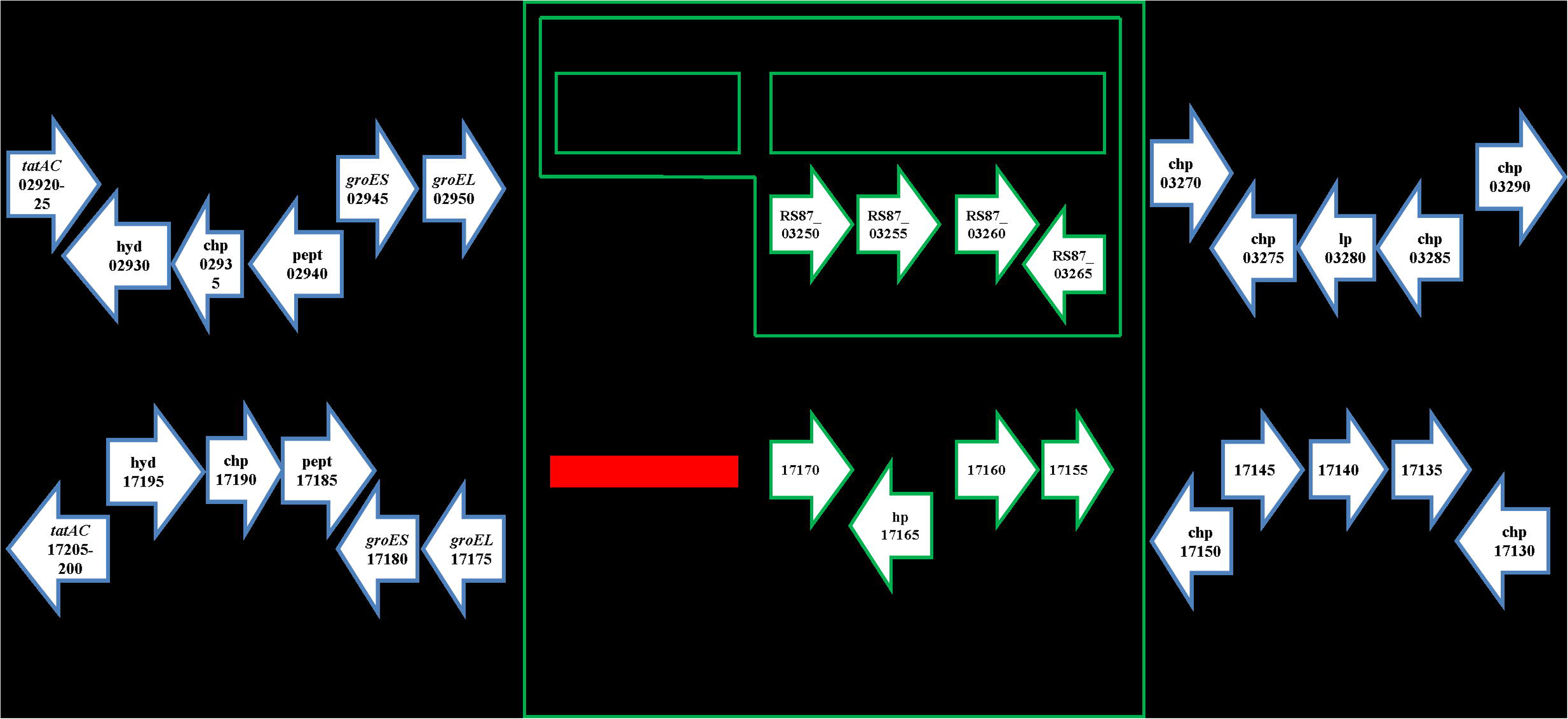
The *Bacillus* bacteriophage SPP1 (NC_004166) homologous region in the *B. safensis* FO-36b genome, as compared with the equivalent genomic region of *B. safensis* JPL_MERTA8-2. Red diamonds denote absence of a single gene/homolog. Red rectangle denotes absence of a series/cluster of ORFs/genes. Green box encloses the phage insertion region.

The *Brevibacillus* phage Jimmer 1 (NC_029104.1) insertion is found to some extent in all 60 draft genomes belonging to the *B. safensis/ B. pumilus* family and the one *Bacillus sp* WP8. In the FO-36b genome, this phage element contains 94 genes (RS87_14155 to RS87_14625). The entire stretch of this insertion can be divided into three blocks, block A (30 genes, RS87_14155 to RS87_14305), block B (30 genes, RS87_14310 to RS87_14455) and block C (34 genes, RS87_14460 to RS87_14625). A major chunk of block C (26 genes RS87_14460 to RS87_14590) is a duplication of block A. The overall scheme of this unique duplication within the insertion is given in Figure 3.

**Figure 3.**
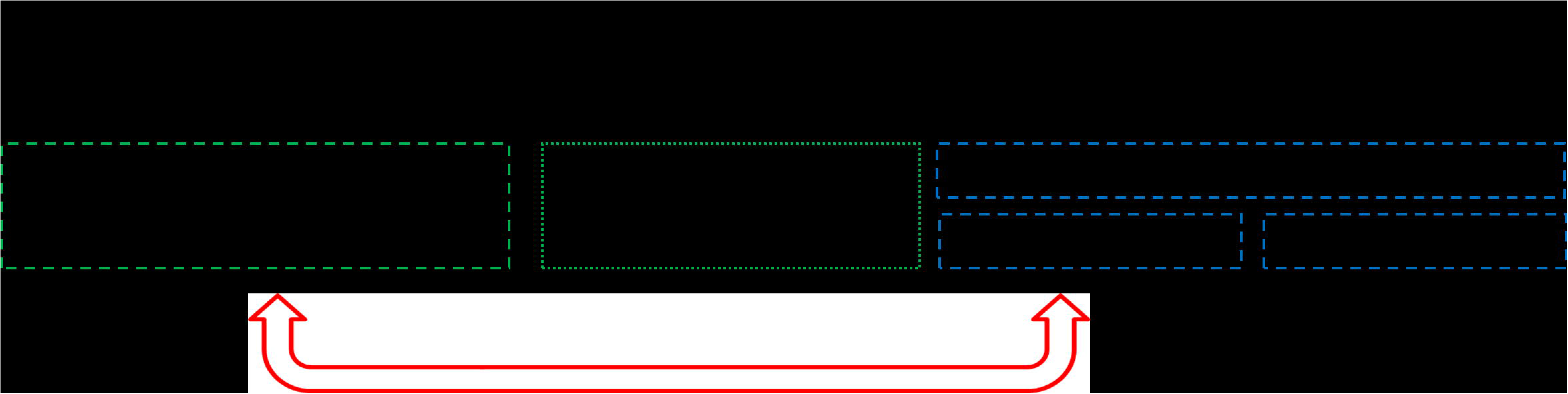
Overall scheme of the *Brevibacillus* phage Jimmer1 (NC_029104) phage insertion in the *B. safensis* FO-36b genome. The three blocks A, B and C and the genes they encompass are shown. The first part of Block C is a duplication of Block A.

A similar version of the Jimmer-1 phage region is found in the non-resistant ATCC7061 (Figure 4). In this case, the block A like region is comprised of 32 ORFs (30 genes and 2 pseudogenes, BAT_0021 to BAT_0052). The block C analog is formed from a cluster of 32 ORFs (29 genes and 3 pseudogenes, BAT_0175 to BAT_0206). Finally, a total of 42 ORFs (41 genes and 1 pseudogene, BAT_0053 to BAT_0094) comprise the equivalent of Block B from FO-36b (Figure 4).

**Figure 4.**
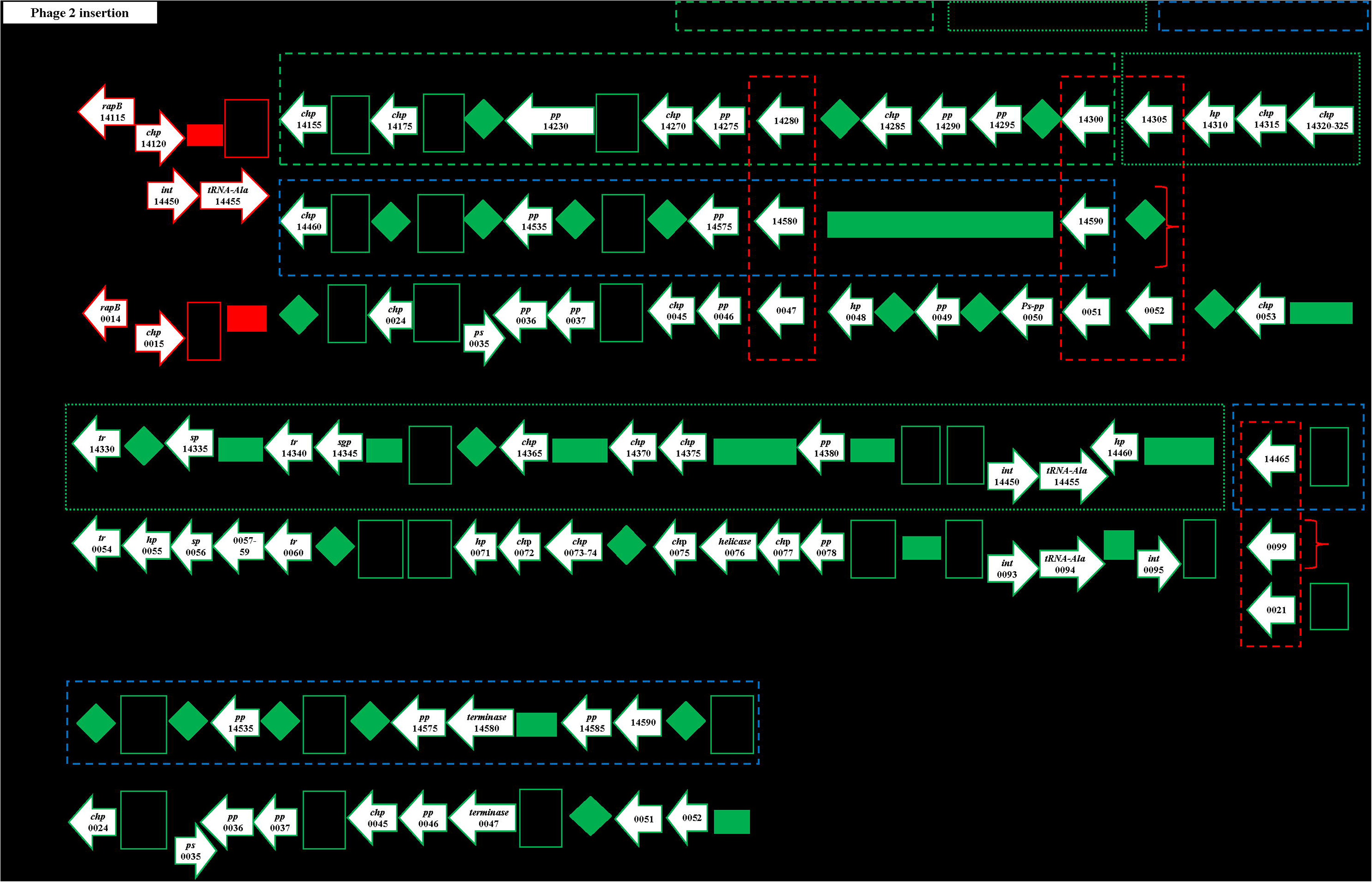
The *Brevibacillus* phage Jimmer1 (NC_029104) phage insertion in the *B. safensis* FO-36b genome as compared with the equivalent region in the genome of *B. pumilus* ATCC7061^T^. Black box encloses the phage insertion region(s). Green (dashed line) box corresponds to block A. Green (dotted line) box corresponds to block B. Blue (dashed line) box corresponds to block C. Red (dashed line) box encloses ‘terminase’ genes. A diamond denotes absence of a single gene/homolog within the phage, while rectangle denotes absence of a cluster of genes/homologs. “hp” = hypothetical protein, “chp” = conserved hypothetical protein, “pp” = phage portal protein, “sp” = structural protein, “sgp” = spore germination protein, “int” = integrase.

The MERTA and SAFR-032 strains show equivalent regions of block A and block C from FO-36b. However, both block B and the duplication of the block A equivalent region are missing in these strains (Figure 5 and Figure 6). The genome of the non-resistant spore producing BSU strain contains the block A and block C equivalents in stretches of 28 ORFs/genes (BSU12810 to BSU12580) and 30 ORFs/genes (BSU12810 to BSU12560) respectively, while block B is entirely missing. However, a major chunk of block A (RS87_14200 to RS87_14300) equivalent region in BSU is duplicated in a stretch of 20 ORFs/genes (BSU25980 to BSU26190) (Figure 7). In general, the occurrence of phage insertion regions and genes therein such as the dUTPase and RecT genes do not appear to be strongly correlated with resistance properties.

**Figure 5.**
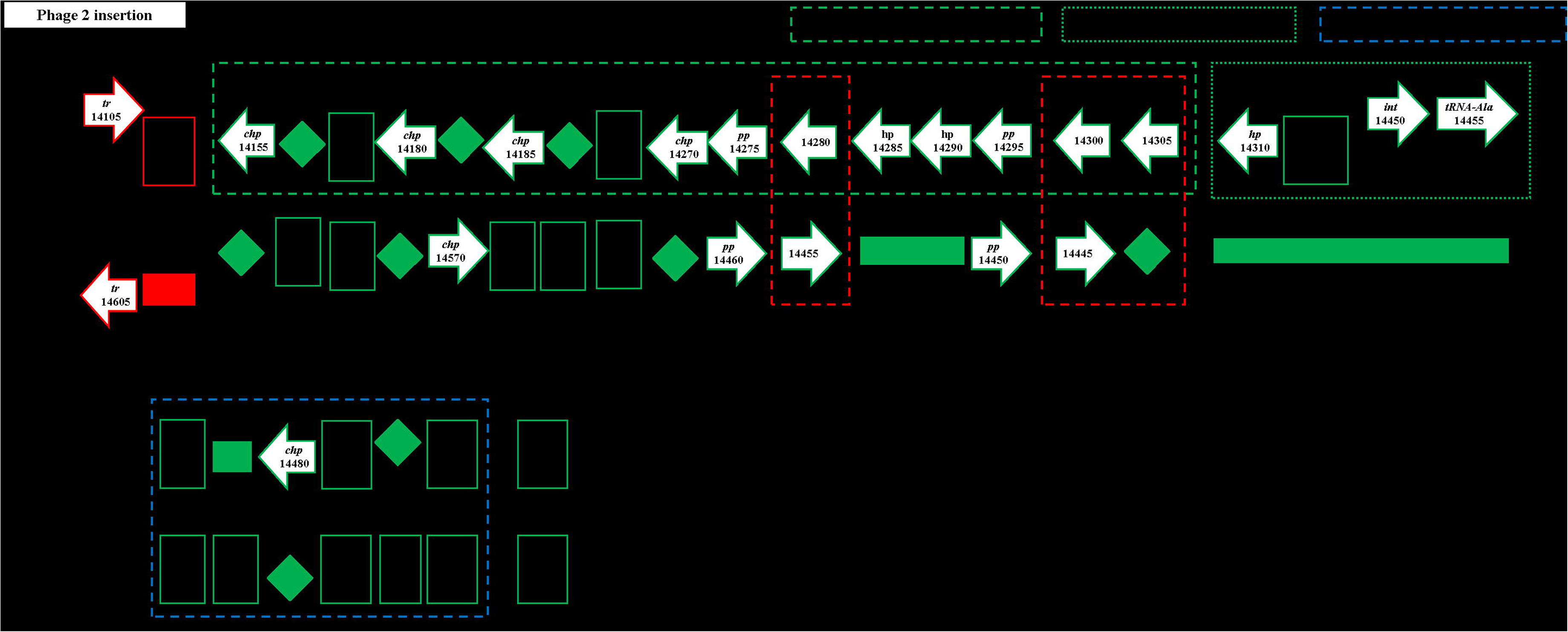
The *Brevibacillus* phage Jimmer1 (NC_029104) phage insertion in the *B. safensis* FO-36b genome as compared with the equivalent region in the genome of *B. safensis* JPL_MERTA8-2. “hp” = hypothetical protein, “chp” = conserved hypothetical protein, “pp” = phage portal protein, “sp” = structural protein, “sgp” = spore germination protein, “int” = integrase.

**Figure 6.**
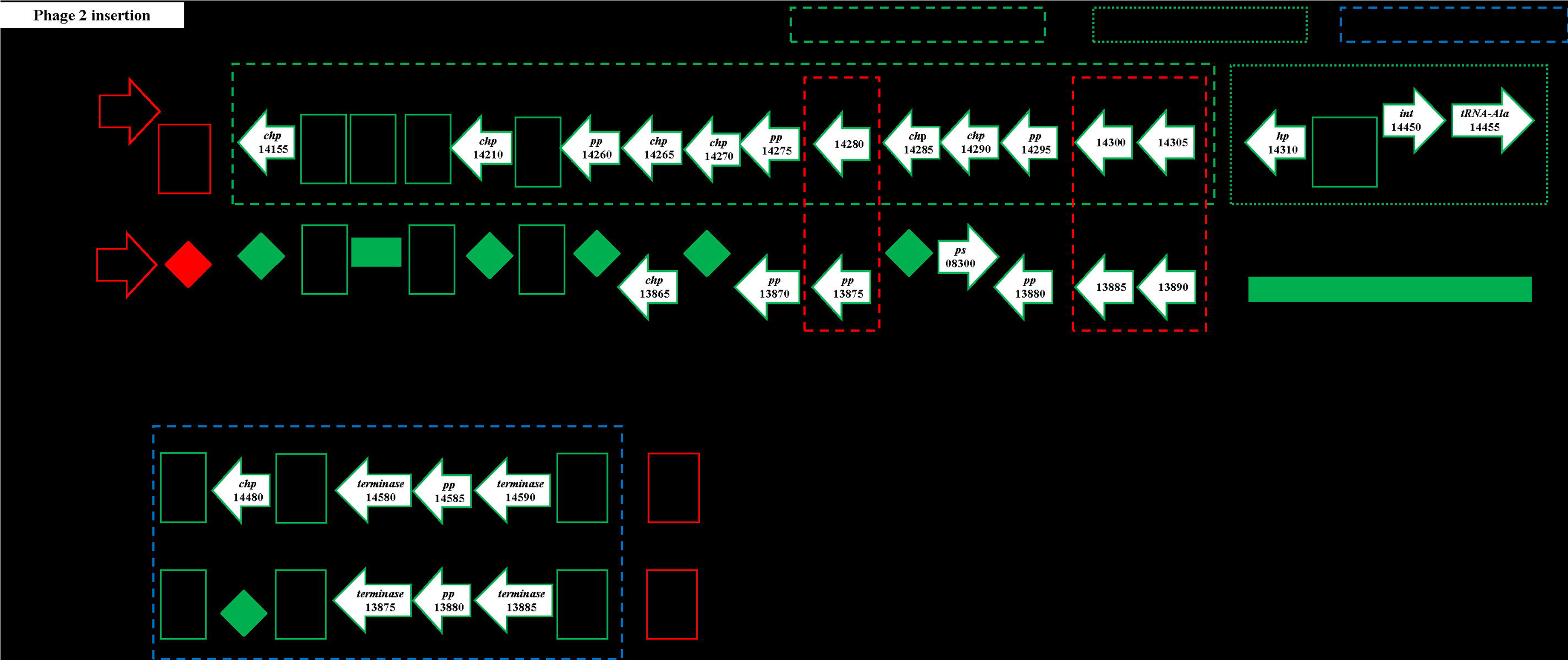
The *Brevibacillus* phage Jimmer1 (NC_029104) phage insertion in the *B. safensis* FO-36b genome as compared with the equivalent region in the genome of *B. pumilus* SAFR-032. “hp” = hypothetical protein, “chp” = conserved hypothetical protein, “pp” = phage portal protein, “sp” = structural protein, “sgp” = spore germination protein, “int” = integrase.

**Figure 7.**
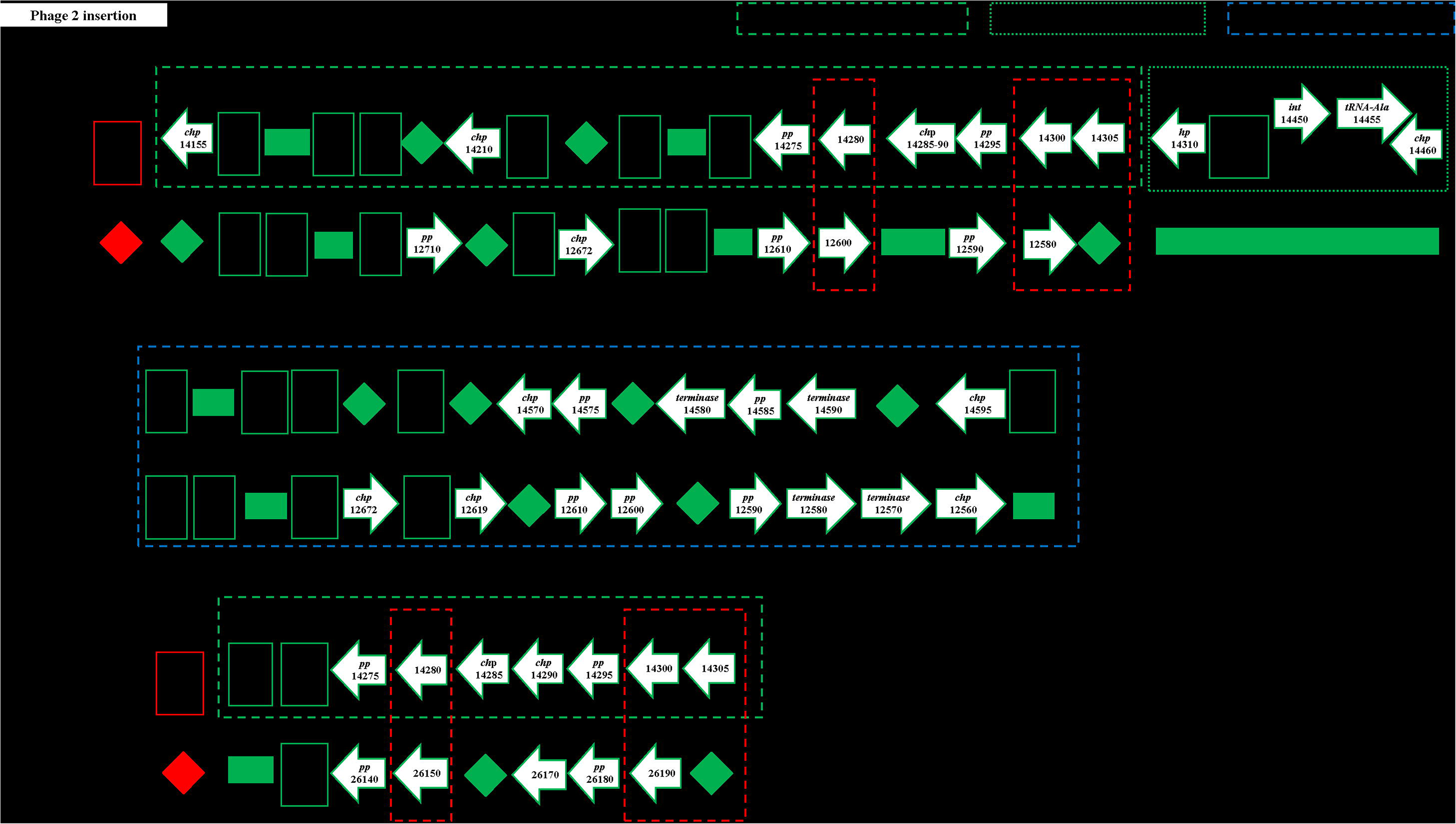
The *Brevibacillus* phage Jimmer1 (NC_029104) phage insertion in the *B. safensis* FO-36b genome as compared with the equivalent region in the genome of *B. subtilis*. “hp” = hypothetical protein, “chp” = conserved hypothetical protein, “pp” = phage portal protein, “sp” = structural protein, “sgp” = spore germination protein, “int” = integrase.

### Genes Shared by FO-36b, SAFR-032, and MERTA but missing in ATCC7061

We had earlier reported that a total of 65 genes that were shared by SAFR-032 and FO-36b, were not found in the ATCC7061 strain [38]. Because they correlate with the presence or absence of resistance, these genes are of potential interest. A re-analysis of this list of genes extending to the MERTA strain showed that 59 of these genes are indeed shared by the MERTA strain as well (Additional file 4: Table S4). All of these genes are shared by at least several of the available 61 *B. pumilus, B. safensis* and *Bacillis sp.* WP8 draft genomes. However, since the resistance properties of these organisms have typically not been examined, it is not immediately possible to determine if the correlation can be extended to these strains.

### Antibiotic Resistance loci in the genomes

The four genomes showed vast differences in the number of antibiotic resistance related mutations that were identified by the CARD [71] search. FO-36b, SAFR-032, MERTA and ATCC7061 had 670, 587, 317, and 495 mutations respectively. BSU comparatively had 861 such mutations. All the four genomes share “*cat86*”, which is a chromosome-encoded variant of the *cat* gene found in *Bacillus pumilus* [72], belonging to the AMR (antimicrobial resistance) gene gamily of chloramphenicol acetyltransferase (CAT).

### Phylogenetic analysis

Previous efforts to define the phylogenetic relationship between various *B. safensis* and *B. pumilus* strains relied on 24 genomes including the unpublished draft sequence (ASJD00000000) of *B. safensis*. Comparing this earlier version with our updated corrected sequence assembly using Mauve shows our version differs considerably (Additional file 5: Figure S1). Given this and the large number of additional draft genomes, it was concluded that a re-analysis would be appropriate. Whole Genome Phylogenetic Analysis and Genome-genome distance analysis were used to examine relationships among the strains. The results of the WGPA are shown in Figure 8, while the GGDC results are given in Additional file 1: Table S1. The phylogenetic trees are consistent with the earlier work (38). Two large clusters are seen. The first consists primarily of strains of *B. pumilus* with no *B. safensis* strains included. The first major cluster is itself broken into two large sub clusters, the first one of which includes both SAFR-032 and ATCC7061. The second sub cluster includes strains from the *B. altitudinis* complex (https://www.ncbi.nlm.nih.gov/Taxonomy/Browser/wwwtax.cgi?id=1792192), as well as other strains recently reported to be *B. pumilus*. The second major cluster consists primarily of *B. safensis* isolates but does include several likely misnamed *B. pumilus* strains too. This latter cluster includes both the FO-36b and the MERTA8-2 strains.

**Figure 8.**
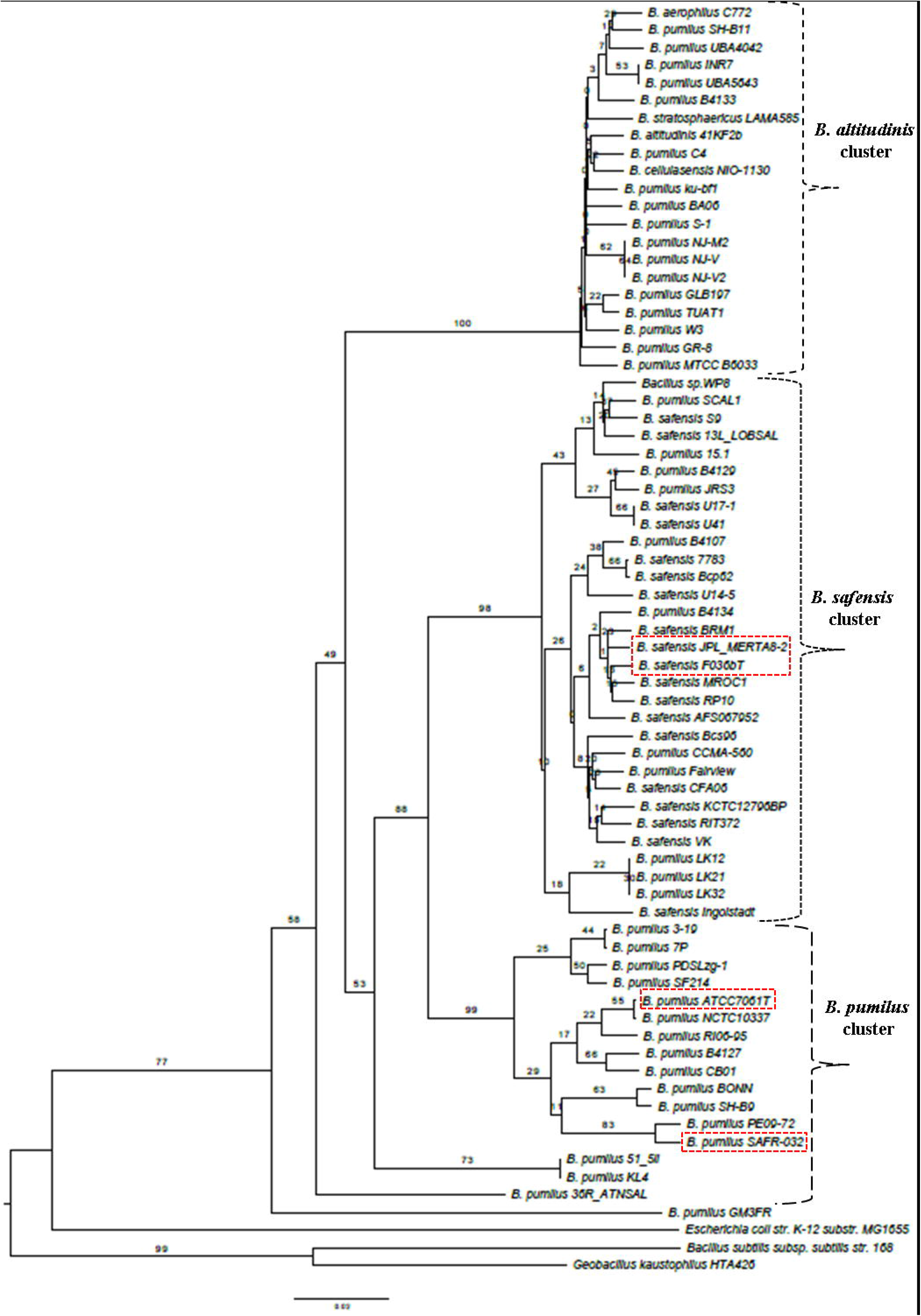
Whole genome Phylogenetic Analysis (WGPA) using the latest version of the Genome-BLAST Distance Phylogeny (GBDP). *B. safensis* FO-36b, *B. safensis* JPL_MERTA8-2B, *B. pumilus* SAFR-032, and *B. pumilus* ATCC7061^T^ are highlighted in red dash-lined rectangles

To further ascertain this observation, a maximum likelihood tree was obtained for the gene *gyrA* (Figure 9), which further supports the WGPA and GGDC analysis. Alternative tree constructions of *gyrA* are provided as Additional file 6: Figure S2 and Additional file 7: Figure S3.

**Figure 9.**
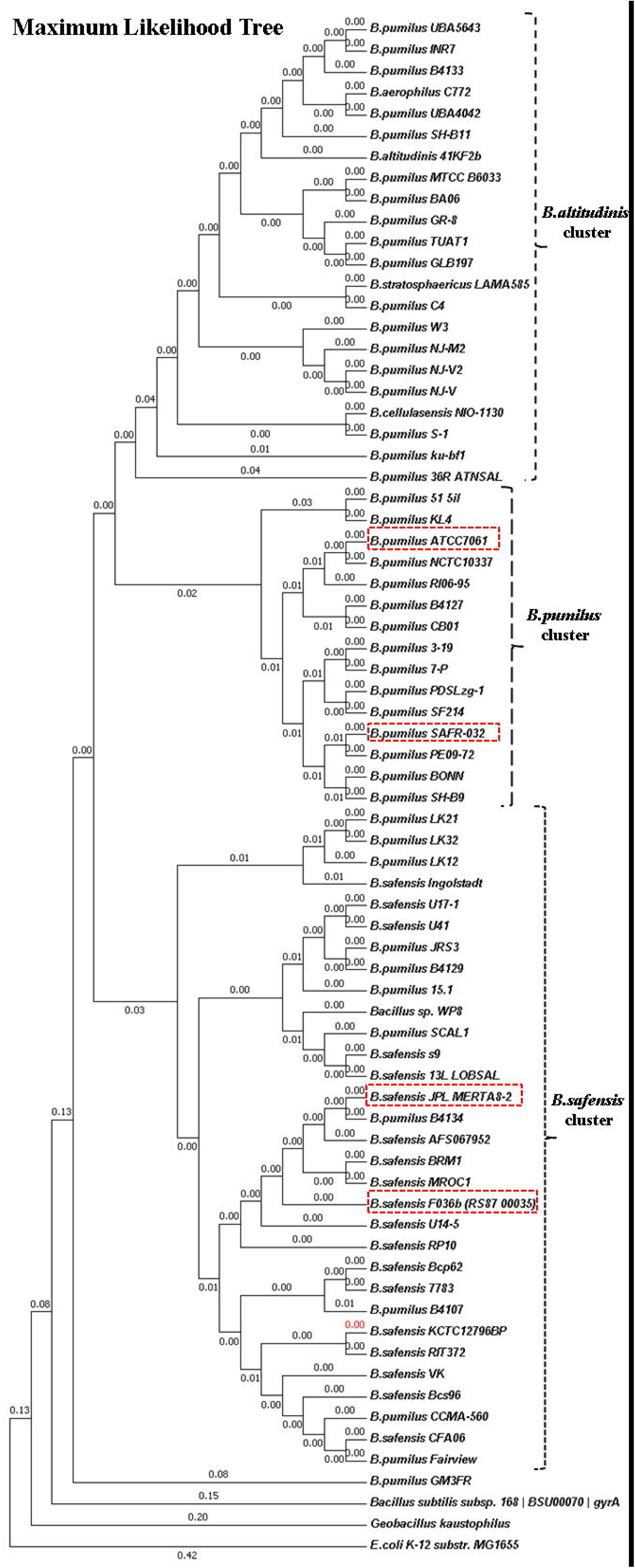
Molecular Phylogenetic analysis by the Maximum Likelihood method. *B. safensis* FO-36b, *B. safensis* JPL_MERTA8-2B, *B. pumilus* SAFR-032, and *B. pumilus* ATCC7061^T^ are highlighted in red dash-lined rectangles.

## Discussion

If there is a single group of genes accounting for the elevated spore resistances seen in various strains of *B. pumilus* and *B. safensis* then the relevant genes should be shared by all three strains examined here but absent in the type strain. The fact that the extent of resistance and type of resistance (radiation, desiccation etc.) varies suggests there may not be a single set of genes involved. In any event, the distinctions in resistance seen may occur due to regulatory differences resulting in key genes associated with resistance being expressed at different levels or for different times [73–75]. Although not correlated with resistance information, it is of interest that in FO-36b, there is a dUTPase and a DNA recombinase gene included in the *Bacillus* bacteriophage SPP1 (NC_004166.2) homologous region.

### Phage insertions

Conjugative elements and phage-mediated insertions play major roles in the evolution of bacteria [76] by contributing to the genetic variability between closely related bacterial strains[77]. Such variability is often implicated in the phenotypical differences such as bacterial pathogenesis [77–80]. Bacteriophage-mediated horizontal gene transfer enhances bacterial adaptive responses to environmental changes such as the rapid spread of antibiotic resistance [81]. Furthermore, phages mediate inversions, deletions and chromosomal rearrangements, which help shunt genes that could directly impact the phenotype between related strains [77] or between phylogenetically distant strains via horizontal gene transfer (HGT)[82]. All of these evolutionary events have implications for selection and fitness.

The first phage insertion in FO-36b is homologous to the *Bacillus* bacteriophage SPP1. The SPP1 is a 44-kb virulent *Bacillus subtilis* phage, well-known for its ability to mediate generalized transduction, a widespread mechanism for the transfer of any gene from one bacterium to another [83]. The second insertion is homologous to *Brevibacillus* phage Jimmer 1, which is one of several myoviruses that specifically target *Paenibacillus larvae*, a *Firmicute* bacterium, as a host [84].

The *B. safensis* strain lacks the ICEBs1-like element that was previously found in SAFR-032 and as an incomplete analog in ATCC7061 [39]. As reported earlier [39], the ICEBs1-like element does harbor some SAFR-032 unique genes and thus, their presence was suggested as being possibly responsible for the resistance properties of SAFR-032. The absence of the ICEBs1-like element in the FO-36b genome suggests that this may not be the case. FO-36b has an established phenotype showing spore resistance to peroxide exceeding that of the other JPL-CRF isolates [13]. SAFR-032 spores have been demonstrated to show resistance to UV radiation exceeding that of the other JPL-CRF isolates [16]. Given that both FO-36b and SAFR-032 harbor genes unique to each of them, on their respective phage elements (the two insertion elements in the case of FO-36b that are reported here and the ICEBs1-like element in the case of SAFR-032), a role of these unique genes in their respective unique spore phenotypes cannot be entirely ruled out.

Furthermore, more than one-half of the *in silico* predicted phage gene products are hypothetical proteins without any assigned functions [85–89]. Comparative genomic approaches use closely related phages from different host organisms and exploit the modular organization of phage genomes [90]. However, these methods are not adequate to address the hypothetical protein coding ORFs that are unique to phage insertions found in a given microbial strain that displays unique phenotypes as in the case of FO-36b and SAFR-032.

Hypothetical phage proteins are considered potential candidates for bacterial detection and antimicrobial target selection. In recent times, efforts towards discovering phage-based antimicrobials have led to the experimental characterization of specific phage proteins [91]. The identification of hypothetical ORFs unique to FO-36b and SAFR-032 phage insertion elements mark them out as potential biomarker candidates for the identification/detection of such strains.

The distribution of the phage elements is not consistently associated with resistance properties. The Jimmer1 phage includes many genes found in all the strains whether resistant or not. The previously highlighted ICEBs1 like element found in the resistant SAFR-032 is not found in the resistant FO-36b strain. The SPP1 element found in the resistant ATCC7061 strain is missing in SAFR-032. One might speculate that individual phage elements might have been transferred to the main genome in the last two cases thereby maintaining consistency with resistance properties. However, no examples of this were found.

### Non-phage associated genes

Genes shared by the three resistant spore producing strains but not the non-resistant ATCC7061 strain are candidates for association with thee resistance properties. Of the 65 ORFs we had reported earlier to be uniquely shared by SAFR-032 and FO-36b [38], 59 are shared by the MERTA strain (Additional file 4: Table S4). When the analysis is extended to all 61 genomes it was found that in each case at least one additional organism had a homolog to the candidate gene. For example, one of these ORFs (FO-36b locus tag RS87_09285), is found to be shared by *B. safensis* MROC1 (isolated from the feces of *Gallus gallus*) and *B. safensis* RP10 (isolated from soils contaminated with heavy metals in Chile). Most of the strains containing these genes are isolates from environments that have some extreme stress component. However, it is not known if the stress component would include resistance to radiation or peroxide. Based on their names alone, some of these strains, such as *B. altitudinis*, and *B. stratosphericus* may be of special interest for further comparison and investigation of their spore resistance properties.

### Highly unique open reading frames

The nine FO-36b ORFs (hypothetical proteins) that were found to be absent from all the *B. safensis/ B. pumilus* (and the *Bacillus sp*. WP8) genomes available in the NCBI database (Table 2A) may be envisioned as possibly contributing to the FO-36b spore resistance. Four of these highly unique ORFs are found on phage elements (one ORF, RS87_03140 on the *Bacillus* bacteriophage SPP1 insertion and three ORFs, viz., RS87_14155, RS87_14285, and RS87_14310 on the *Brevibacillus* phage Jimmer 1 insertion). This is similar to the situation with the ICEBs-1 like element in SAFR-032 that harbors unique SAFR-032 ORFs [39]. Four other ORFs had fewer than 5 homologs found in other *B. pumilus*/*B. safensis* genomes. Two of these four ORFs, are also found on the phage elements and hence could be random remnants of lateral transfer.

### Genes involved in peroxide resistance and DNA repair

We had previously reported 15 peroxide resistance genes in SAFR-032, of which 2 were not shared by either the earlier draft version of FO-36b, or the type strain ATCC7061 [38]. Five of these peroxide genes were uniquely shared by SAFR-032 and the earlier draft version of the FO-36b genome. Of the 8 SAFR-032 DNA repair genes reported then, 5 were not shared by FO-36b or ATCC7061. We verified those results against the now complete FO-36b genome, and the status of the genes remains the same as before.

We also looked at the gene coding for ‘Dps’, which is a DNA-binding protein. Dps is very well-characterized for providing protection to cells during exposure to severe environmental conditions such as oxidative stress and nutritional deprivation in gram negative bacteria such as *E. coli* [92] as well as gram positive *Firmicutes* species such as *Staphylococcus aureus* [93], *B. subtilis* [94], *B. anthracis* [95, 96] and *B. cereus* [97, 98]. With its tripartite involvement in DNA binding, iron sequestration, and ferroxidase activity, Dps plays important roles in iron and hydrogen peroxide detoxification and acid resistance [99, 100]. The homolog for the *dps* gene in *Bacillus* strains is ‘*mrgA’* [101], which is highly conserved amongst the resistant spore-producing FO-36b and SAFR-032, as well as the non-resistant spore-producing ATCC7061 strain. Likewise, other peroxide resistance genes were checked for their presence/absence and were all found conserved in the four genomes. Thus it is unlikely that any of these genes play any role in the resistances seen in *B. safensis* FO-36b and *B. pumilus* SAFR-032.

### Antibiotic resistance

There is increasing concern about bacterial pathogenicity under microgravity and/or in human spaceflight [102]. This is validated by reports that several microbial strains isolated from, or exposed to space environments, show resistance to desiccation, heat-shock, and/or applied antibiotics [103, 104]. A global analysis of the four genomes was undertaken to identify the presence of known antibiotic resistance related mutations. It was found that the FO-36b and SAFR genomes had significantly larger numbers (approximately 100-200 more) of the mutations as compared with the MERTA and ATCC7061 genomes. On a comparative scale, the genome of BSU had almost 200 more AMR related mutations. The mere presence or the number of these mutations as such cannot be linked with the respective antibiotic resistance properties of these strains. However, further analysis of antibiotic susceptibility of these strains is warranted to establish how they differ from other strains.

### Phylogenetic analysis

The current study used Whole Genome Phylogenetic Analysis methodology to delineate phylogenetic distances based on whole genomes of organisms. This and the separate genome-genome distance analysis are consistent with, but more detailed than the earlier study [38]. Additionally, the “*gyrA*” tree analysis was found to support the WGPA and GGDC results. In agreement with the earlier studies, the *B. safensi*s/ *B. pumilus* strains form a coherent cluster with three large sub clusters (Figure 8, 9). One of the large sub clusters includes the FO-36b, and MERTA strains as well as all other *B. safensis* strains. *B. pumilus* strains in this grouping may be usefully renamed as *B. safensis*. SAFR-032 and ATCC7061 are in a second sub cluster that is exclusively populated with *B. pumilus* strains. The third sub cluster includes all members of the *B. altitudinis* group and many *B. pumilus* strains.

## Conclusions

A recent report [105] has implicated that the opposing effects of environmental DNA damage and DNA repair result in elevated rates of genome rearrangements in radiation-resistant bacteria that belong to multiple, phylogenetically independent groups including *Deinococcus*. This view is not consistent with the four genomes examined in detail here as few arrangements are observed. Comparison with earlier results [38, 39] did not yield anything new and thus although candidates continue to exist, no specific gene has been identified as likely being responsible for the resistances exhibited by these organisms. The differences in resistance properties can easily be attributed to changes in expression level but of what gene or genes? With a larger phylogenetic tree now available, it should be possible to select a representative subset of strains for further resistance studies as well as sequencing.

## List of abbreviations used

FO-36b: *B. safensis* FO-36b^T^ (Genbank Accession no: - CP010405).
SAFR-032: *B. pumilus* SAFR-032.
ATCC7061: *B. pumilus* ATCC7061^T^.
MERTA: *B. safensis* JPL-MERTA-8-2.
BSU: *B. subtilis subsp. subtilis str.* 168.

## Declarations

### 1. Ethics approval and consent to participate

Not Applicable

### 2. Consent for publication

Not applicable

### 3. Availability of data and material

The datasets used and analyzed within the current study are available from the NCBI Website as referenced in the paper. The sequence of the *B. safensis* FO-36b strain is being deposited with the NCBI/Genbank with the accession number CP010405. Until the deposit is complete, the data will be available from the corresponding author.

### 4. Competing Interests

All authors declare they have no competing interests.

### 5. Funding

This work was funded in part by NASA Grant NNX14AK36G to GEF. Discussions relating to the NASA project led to this work and provided resources for the design of the study, and writing of the manuscript.

### 6. Author’s contributions

KJV, VGS, and GEF conceived and designed the study. MRT annotated and curated the annotated genome, analyzed the data, performed the comparative genome analysis. SM and MRT prepared the tables and the figures. VGS prepared the library for NextGen sequencing and processed the resulting data. AW performed the sequencing. VGS and ROG conducted local sequencing studies to order contigs and close the genome. KRG provided genomic DNA. MRT, VGS and GEF prepared a draft paper which was finalized with help from all the authors. All authors read and approved the final manuscript.

## 7. Acknowledgements

The authors acknowledge the use of the Maxwell/Opuntia Cluster and the advanced support from the Center of Advanced Computing and Data Systems at the University of Houston to carry out the research presented here. The authors would also like to thank Dr. Jan Meier-Kolthoff, Department of Bioinformatics, Leibniz Institute DSMZ (German Collection of Microorganisms and Cell Cultures), Inhoffenstraße 7 B, 38124 Braunschweig, Germany, for help with the Whole Genome Phylogenetic Analysis. This work was supported in part by NASA Grant NNX14AK36G to GEF.

## Endnotes

None

## Additional files

**Additional file 1: Table S1**. In silico DNA-DNA hybridization (DDH) values showing Genome-genome distance [50] relationship values for the genomes of various *B. pumilus, B. safensis, B. altitudinis* strains. The genomes of *Geobacillus kaustophilus, and B. subtilis subsp. subtilis str.* 168 serving as outliers in the *Firmicutes* group and that of gram-negative *E.coli* MG1655, as a non-*Firmicutes* outlier.

**Additional file 2: Table S2**. Presence and absence of the *B. safensis* FO-36b CRISPR module element protein(s) in the other *B. pumilus* / *B. safensis* genomes.

**Additional file 3: Table S3**. *B. safensis* FO-36b characteristic genes (ORFs/genes that are absent from *B. pumilus* SAFR-032, *B. pumilus* ATCC7061^T^, and, *B. safensis* JPL-MERTA-8-2) and their occurrence (presence/absence) in the *B. pumilus*/*B. safensis* genomes available in the NCBI database. P: Present, A: Absent, *found on phage insertions.

**Additional file 4: Table S4.** *B. safensis* F0-36b genes reported earlier as shared by *B. pumilus* SAFR-032 and not found in the *B. pumilus* ATCC7061^T^ strain [38], compared with the *B. safensis* JPL-MERTA-8-2 strain, and the other *B. pumilus / B. safensis* genomes.

**Additional file 5: Figure S1**. Whole genome alignment of the previously existing *B. safensis* FO-36b sequence (GCA_000691165.1 / ASJD00000000) with our current updated sequence (CP010405) using Mauve [70].

**Additional file 6: Figure S2.** Molecular Phylogenetic analysis by the Neighbor-Joining method. *B. safensis* FO-36b, *B. safensis* JPL_MERTA8-2B, *B. pumilus* SAFR-032, and *B. pumilus* ATCC7061^T^ are highlighted in red dash-lined rectangles.

**Additional file 7: Figure S3.** Molecular Phylogenetic analysis using the Minimum Evolution method. *B. safensis* FO-36b, *B. safensis* JPL_MERTA8-2B, *B. pumilus* SAFR-032, and *B. pumilus* ATCC7061^T^ are highlighted in red dash-lined rectangles.

